# Comparative analysis of antiviral efficacy of FDA-approved drugs against SARS-CoV-2 in human lung cells: Nafamostat is the most potent antiviral drug candidate

**DOI:** 10.1101/2020.05.12.090035

**Authors:** Meehyun Ko, Sangeun Jeon, Wang-Shick Ryu, Seungtaek Kim

## Abstract

Drug repositioning represents an effective way to control the current COVID-19 pandemic. Previously, we identified 24 FDA-approved drugs which exhibited substantial antiviral effect against SARS-CoV-2 in Vero cells. Since antiviral efficacy could be altered in different cell lines, we developed an antiviral screening assay with human lung cells, which is more appropriate than Vero cell. Comparative analysis of antiviral activities revealed that nafamostat is the most potent drug in human lung cells (IC_50_ = 0.0022µM).

---

COVID-19 is an emerging infectious disease caused by a coronavirus (1). The causative virus was named as SARS-CoV-2 because it is very similar to SARS-CoV (79.5%) and this virus belongs to the *Betacoronavirus* genus within the *Coronaviridae* family (2). Both SARS- and MERS-CoV also belong to the same *Betacoronavirus* genus.

Neither vaccine nor therapeutic has been developed for SARS- and MERS-CoV and the current standard of care for the patients with COVID-19 is just supportive care. However, numerous clinical trials are ongoing globally with FDA-approved drugs as drug repositioning programs (https://www.covid-trials.org/). Among these drugs, (hydroxy)chloroquine, lopinavir/ritonavir, and remdesivir are those that are the most frequently being tested worldwide due to the well-known in vitro antiviral effects on both MERS- and SARS-CoV and even on SARS-CoV-2 (3).

In our previous drug repositioning study, we identified a total of 24 potential antiviral drug candidates from FDA-approved drugs (4). These drugs showed very potent antiviral efficacy against SARS-CoV-2 (0.1 µM < IC_50_ < 10 µM) in the experiments using Vero cells. Although Vero cells are commonly used for virus infection and propagation, they were originally isolated from the African green monkey kidney, thus do not represent the respiratory cells from the human lung, which is the main target tissue for SARS-CoV-2 infection. In this study, we compared the antiviral efficacy of the 24 potential antiviral drug candidates against SARS-CoV-2 using Calu-3 human lung cells. Calu-3 was originally isolated from human lung adenocarcinoma and is a well-characterized epithelial cell line (5).

In order to conduct the dose-response curve (DRC) analysis with drugs, Calu-3 cells were treated with each drug candidate 24 h prior to SARS-CoV-2 infection. The infected cells were incubated for another 24 h and then fixed for immunofluorescence. Both viral N protein and host cell nucleus were stained by immunofluorescence and the quantitative analysis to measure the inhibition of virus infection and the cell viability due to drug treatment was conducted using our in-house image mining (IM) software.

The DRC analysis of the reference drugs (i.e., chloroquine, lopinavir, and remdesivir) (Fig. 1) showed differences in IC_50_ in between Vero and Calu-3 cells. While the IC_50_ values of both chloroquine and lopinavir increased by ∼ 10 fold and ∼ 2 fold, respectively (Table 1), the IC_50_ of remdesivir rather decreased by 10 fold compared to that with Vero cells, perhaps due to the low metabolic capacity in Vero cells (6)(Table 2). These discrepancies might in part account for the different outcomes from numerous clinical trials using chloroquine, lopinavir, and remdesivir. So far, the treatment with (hydroxy)chloroquine or lopinavir/ritonavir did not show any promising results concerning the COVID-19 treatment (7)(8); however remdesivir seems to be effective for treatment of patients with COVID-19 in certain clinical settings (https://www.nbcnews.com/health/health-news/coronavirus-drug-remdesivir-shows-promise-large-trial-n1195171).

**Table 1.** List of drugs with increased IC_50_ in Calu-3 cells

| Drug name | IC<sub>50</sub> in Vero (μM)<sup>a</sup> | IC<sub>50</sub> in Calu-3 (μM)<sup>b</sup> | Fold change |
| --- | --- | --- | --- |
| Fold change > 4 |  |  |  |
| Tetrandrine | 3 | 13.5 | 4.50 |
| Berbamine hydrochloride | 7.87 | >50 | 6.35 |
| Abemaciclib | 6.62 | 43.7 | 6.60 |
| Cepharanthine | 4.47 | 30 | 6.71 |
| Gilteritinib | 6.76 | >50 | 7.40 |
| Chloroquine | 7.28 | 69.2 | 9.51 |
| Amodiaquine dihydrochloride | 5.15 | >50 | 9.71 |
| Mefloquine | 4.33 | >50 | 11.55 |
| Fold change > 2 |  |  |  |
| Salinomycin sodium | 0.24 | 0.5 | 2.08 |
| Lopinavir | 9.12 | 21.7 | 2.38 |
| Ciclesonide | 4.33 | 10.64 | 2.46 |
| Proscillaridin | 2.04 | 5.95 | 2.92 |
| Niclosamide | 0.28 | 0.84 | 3.00 |
| Anidulafungin | 4.64 | 17.23 | 3.71 |
| Digoxin | 0.19 | 0.72 | 3.79 |
| Bazedoxifene | 3.44 | 12.63 | 3.67 |

<sup>a</sup> IC<sub>50</sub> was determined by Jeon et al. (4).
<sup>b</sup> IC<sub>50</sub> was determined in this study.

**Table 2.** List of drugs with decreased IC_50_ in Calu-3 cells

| Drug name | IC<sub>50</sub> in Vero (μM)<sup>a</sup> | IC<sub>50</sub> in Calu-3 (μM)<sup>b</sup> | Fold change |
| --- | --- | --- | --- |
| Nafamostat mesylate | 13.88 | 0.0022 | 0.00016 |
| Remdesivir | 11.41 | 1.3 | 0.11 |
| Hydroxyprogesterone caproate | 6.3 | 3.87 | 0.61 |
| Digitoxin | 0.23 | 0.16 | 0.70 |
| Cyclosporine | 5.82 | 4.69 | 0.81 |

<sup>a</sup> IC<sub>50</sub> was determined by Jeon et al. (4) except for nafamostat mesylate.
<sup>b</sup> IC<sub>50</sub> was determined in this study.

**Figure 1.**
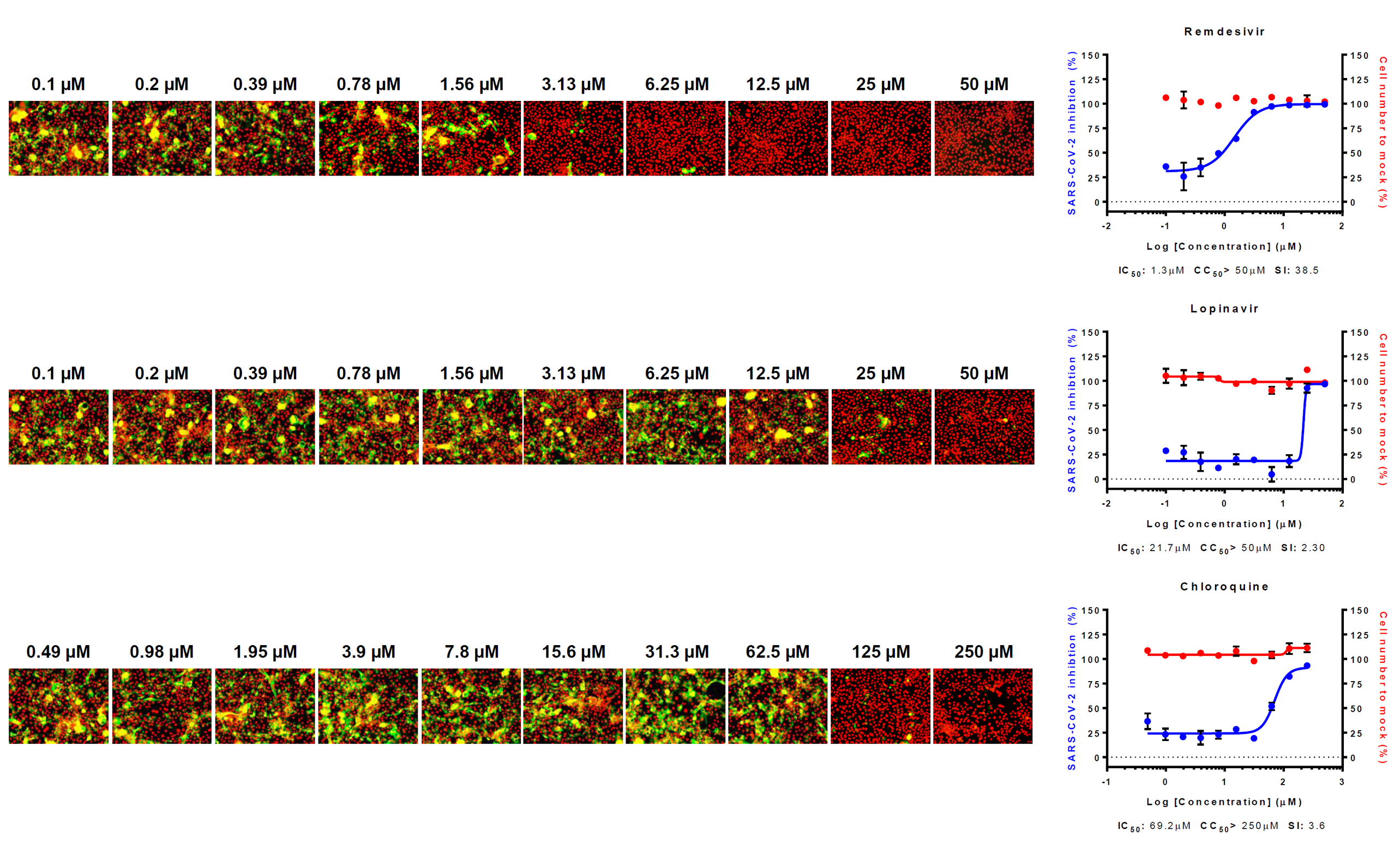
Dose-response curve (DRC) and images of reference drugs. Three reference drugs - remdesivir, lopinavir and chloroquine – were serially diluted by 2-fold to generate a 10-point DRC. Graphs are shown on the right and representative images of each point are shown in the left. Red line indicates percentage of cell number compared to mock infection control, and blue line indicates percent infectivity compared to DMSO control. SARS-CoV-2 infectivity was measured by immunofluorescence of SARS-CoV-2 N protein. Each point is a mean of duplicate experiments ± standard deviation (SD). IC_50_, CC_50_ and selective index (SI) is noted below each graphs. Nucleus is shown in red, and viral N protein is shown in green.

Interestingly, the IC_50_ values of most drugs in our study increased in varying degrees in Calu-3 cells (Fig. 2A and C) (Table 1 and 3). Only 5 drugs showed decreases in IC_50_ (Fig. 2B) (Table 2): Nafamostat mesylate, Remdesivir, Hydroxyprogesterone caproate, Digitoxin, Cyclosporine. Although nafamostat mesylate was not included in our earlier study, we compared the antiviral efficacy of this drug at this time in between Vero and Calu-3 cells following the discovery that TMPRSS2, a host protease necessary for priming viral spike glycoprotein, could be a target for COVID-19 antiviral development (9). The discrepancy in IC_50_ was remarkable with nafamostat mesylate; the IC_50_ decreased by ∼ 6,000 fold when the drug was used in the SARS-CoV-2-infected Calu-3 cells perhaps due to the dominant role of TMPRSS2-dependent viral entry in the Calu-3 human lung epithelial cells (10)(11). In addition, the IC_50_ of nafamostat mesylate was exceptionally low (0.0022 µM), which indicates that nafamostat mesylate is ∼ 600-fold more potent than remdesivir in Calu-3 cells. It became more apparent that blood clotting is one of the complicating manifestations in patients with COVID-19 (12)(13), and nafamostat mesylate may play dual roles not only as an antiviral to block viral entry but also as an anticoagulant to remove blood clots frequently associated with acute respiratory distress syndrome (ARDS).

**Table 3.** List of drugs with unchanged IC_50_ in Calu-3 cells

| Drug name | IC<sub>50</sub> in Vero (μM)<sup>a</sup> | IC<sub>50</sub> in Calu-3 (μM)<sup>b</sup> | Fold change |
| --- | --- | --- | --- |
| Ouabain | <0.1 | <0.1 | 1.00 |
| Eltrombopag | 8.27 | 8.38 | 1.01 |
| Loperamide hydrochloride | 9.27 | 12.53 | 1.35 |
| Hexachlorophene | 0.9 | 1.48 | 1.64 |
| Ivacaftor | 6.57 | 11.55 | 1.76 |
| Oxyclozanide | 3.71 | 6.78 | 1.83 |

<sup>a</sup> IC<sub>50</sub> was determined by Jeon et al. (4).
<sup>b</sup> IC<sub>50</sub> was determined in this study.

**Figure 2.**
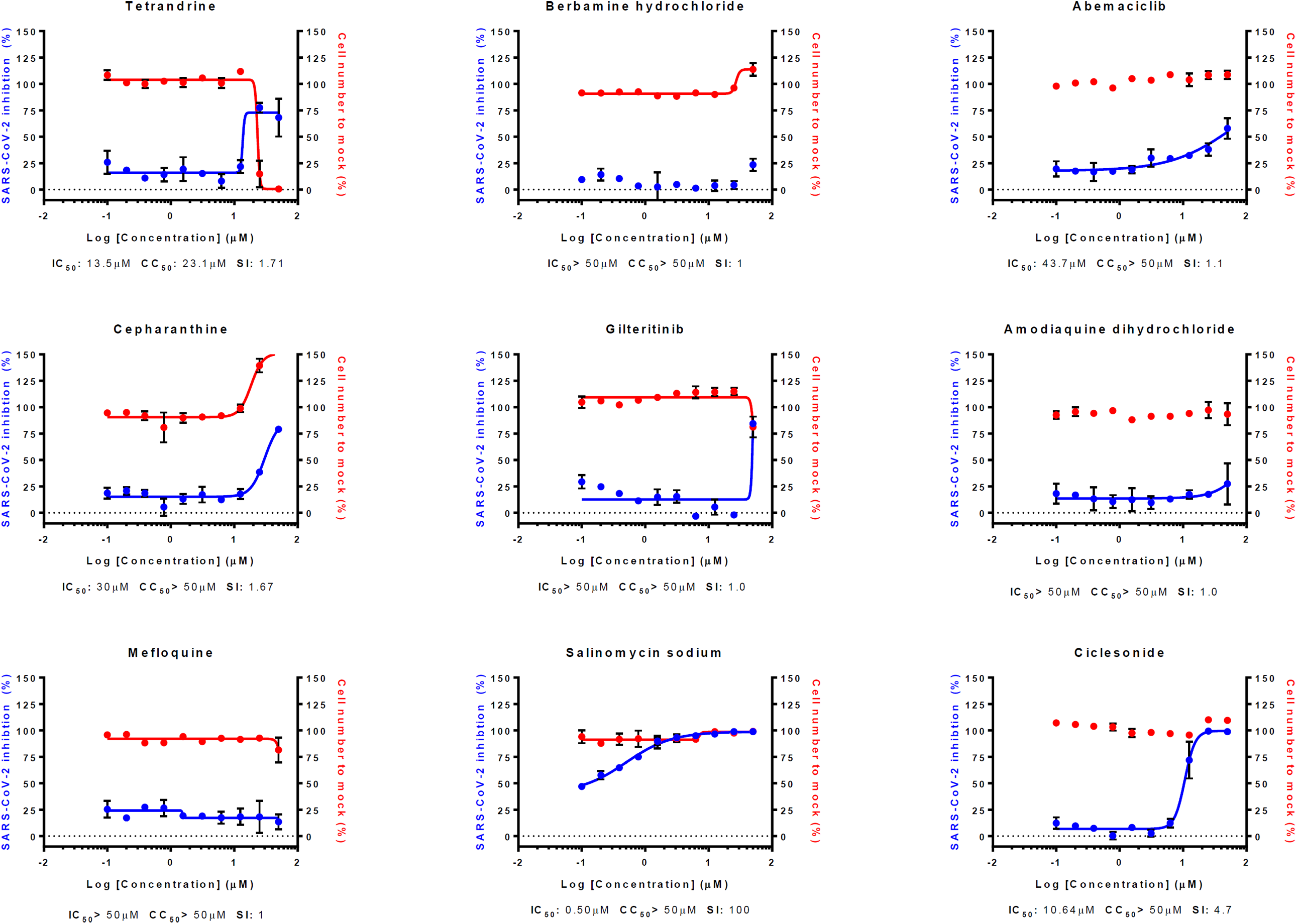
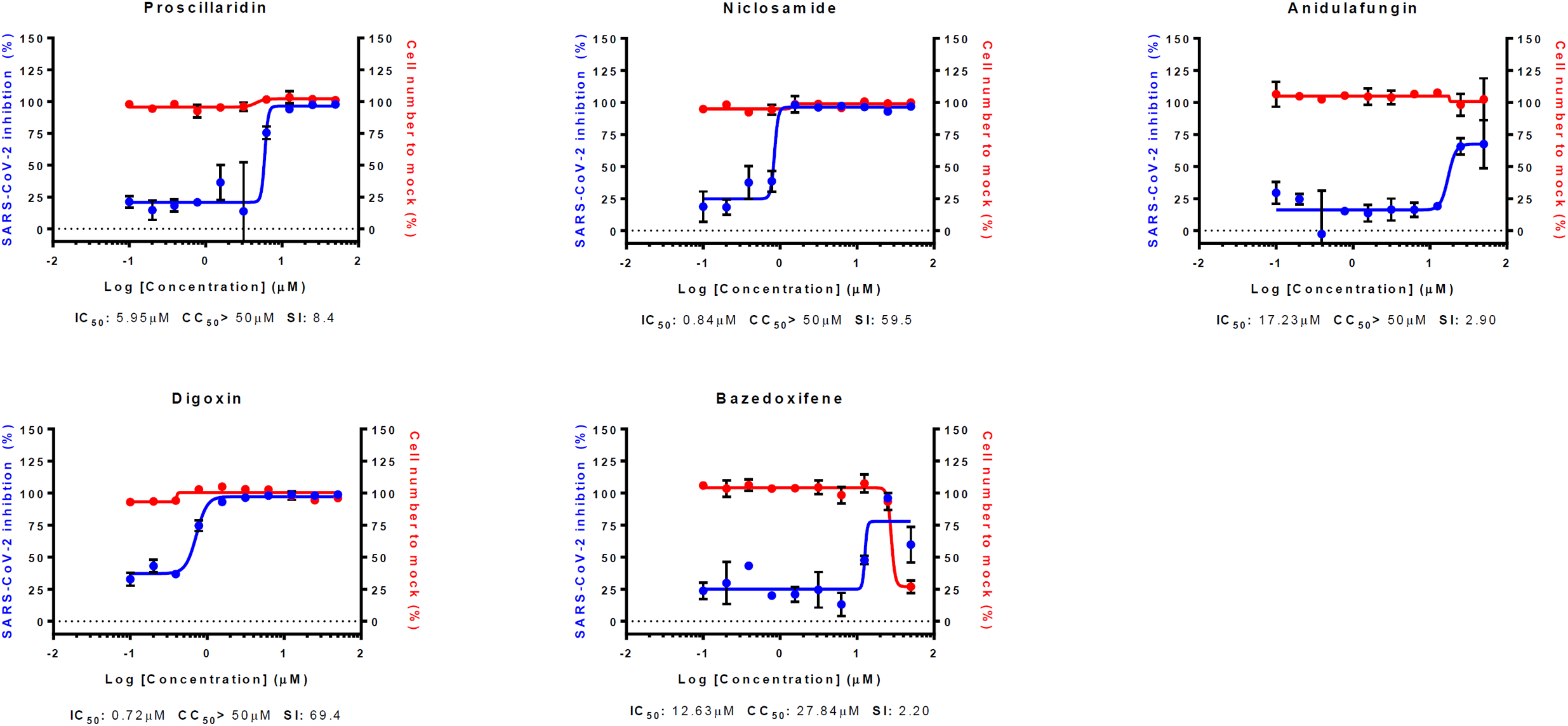
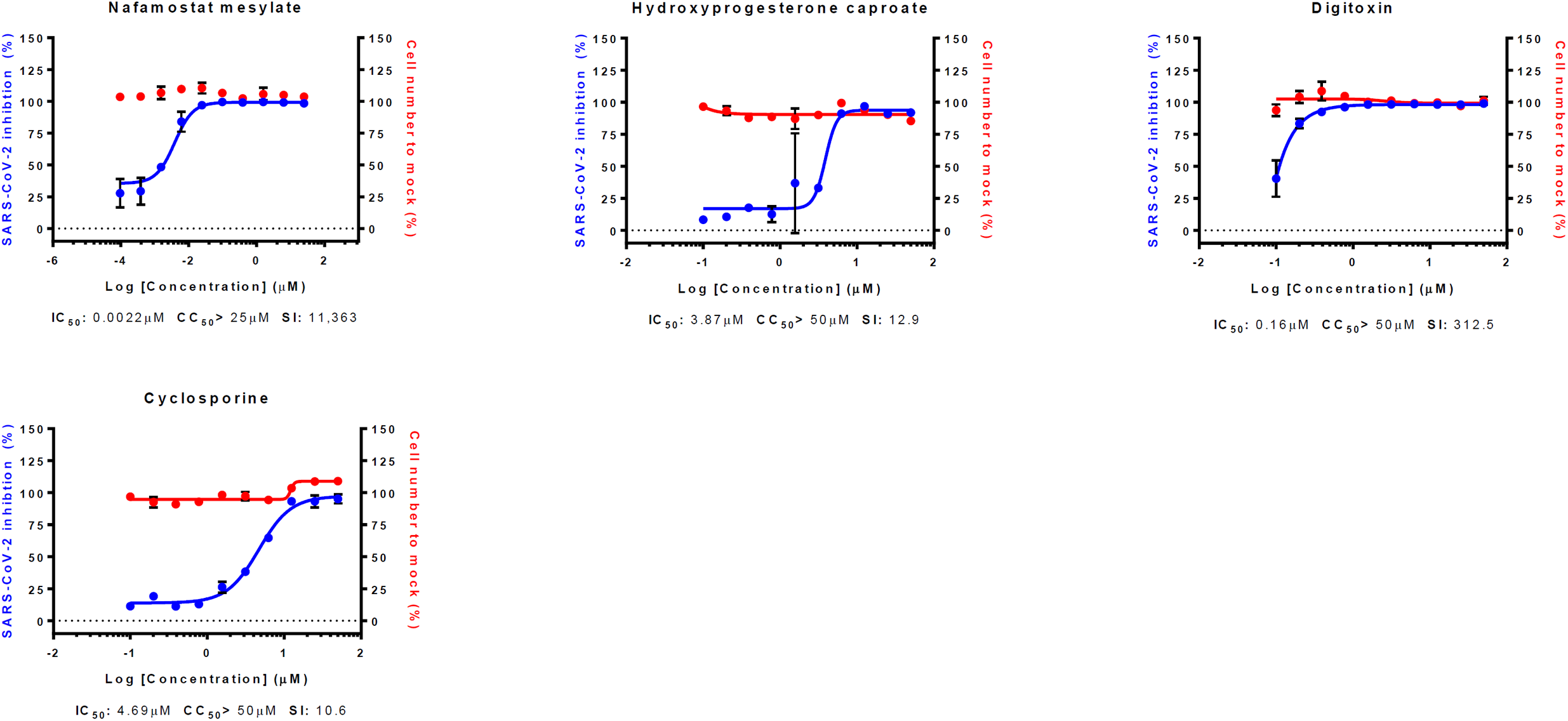
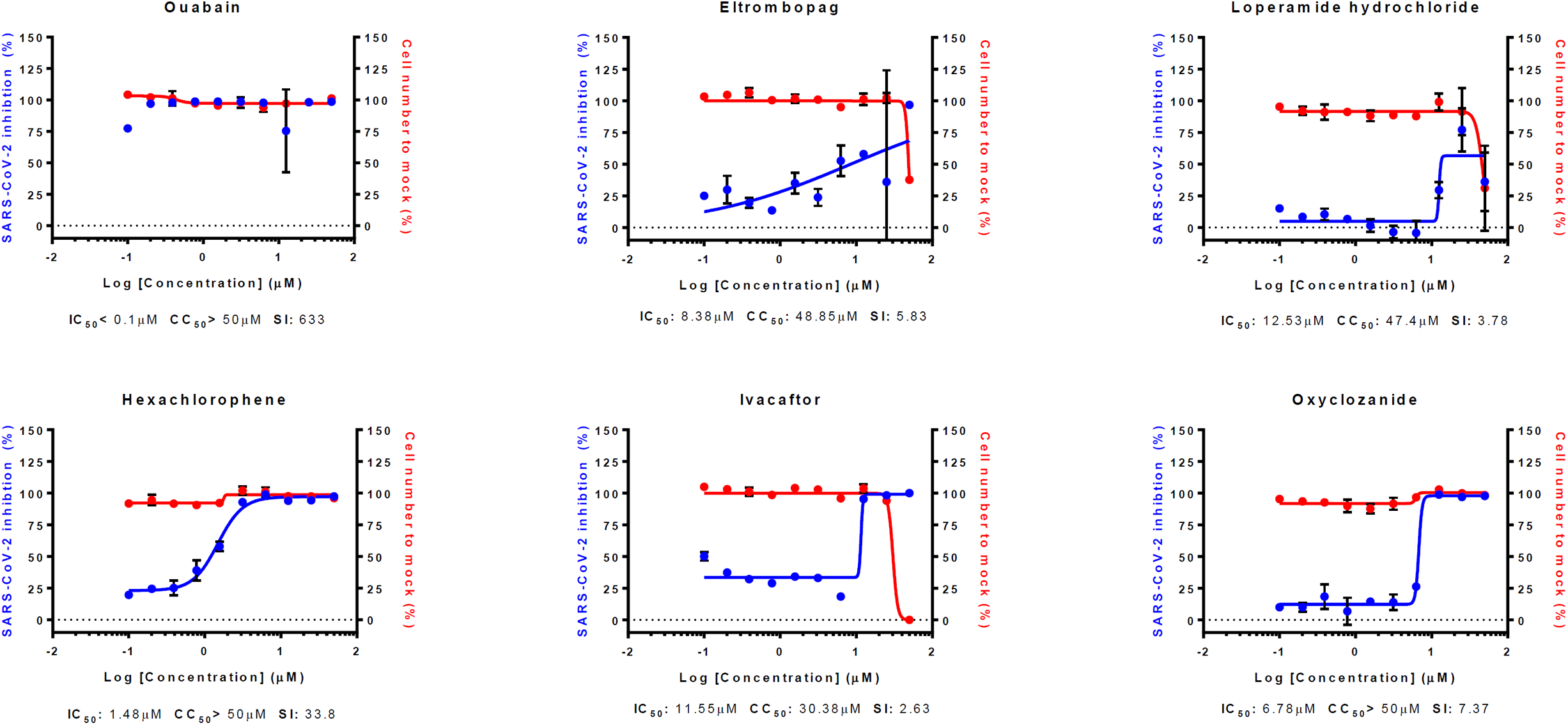
DRC of 24 drugs in Calu-3 that were previously reported in Vero cells in addition to nafamostat mesylate. (A) DRC of compounds with increased IC_50_ value compared to Vero cells (fold change above 2). (B) DRC of compounds with decreased IC_50_ value compared to Vero cells (fold change less than 1). (C) DRC of compounds with unchanged IC_50_ value compared to Vero cells (fold change approximately 1). Red line indicates percentage of cell number compared to mock infection control, and blue line indicates percent infectivity compared to DMSO control. SARS-CoV-2 infectivity was measured by immunofluorescence of SARS-CoV-2 N protein. Each point is a mean of duplicate experiments ± standard deviation (SD). IC_50_, CC_50_ and selective index (SI) is noted below each graphs.

In summary, we compared antiviral efficacy of the potential antiviral drug candidates against SARS-CoV-2 in between Vero and Calu-3 cells and found that nafamostat mesylate is the most potent antiviral drug candidate in vitro. Importantly, nafamostat mesylate has been approved for human use in Japan and Korea for over a decade, thus it can be readily repurposed for COVID-19 following phase II-III clinical trials. Currently, a few clinical trials has been registered (https://clinicaltrials.gov/). According to our results, although in vivo animal models are preferred experimental systems for evaluating antiviral efficacy, in vitro testing using human lung cells is a viable option in addition to the commonly used Vero or VeroE6 cells for assessment of antiviral efficacy when the animal models are not readily available.

## Materials and Methods

### Virus and Cells

Calu-3 used in this study is a clonal isolate, which shows higher growth rate compared to the parental Calu-3 obtained from the American Type Culture Collection (ATCC CCL-81). Calu-3 was maintained at 37°C with 5% CO_2_ in Eagle’s Minimum Essential Medium (EMEM, ATCC), supplemented with 10% heat-inactivated fetal bovine serum (FBS) and 1X Antibiotic-Antimycotic solution (Gibco). SARS-CoV-2 (βCoV/KOR/KCDC03/2020) was provided by Korea Centers for Disease Control and Prevention (KCDC), and was propagated in Vero cells. Viral titers were determined by plaque assays in Vero cells. All experiments using SARS-CoV-2 were performed at Institut Pasteur Korea in compliance with the guidelines of the KNIH, using enhanced Biosafety Level 3 (BSL-3) containment procedures in laboratories approved for use by the KCDC.

### Reagents

Chloroquine diphosphate (CQ; C6628) was purchased from Sigma-Aldrich (St. Louis, MO), lopinavir (LPV; S1380) was purchased from SelleckChem (Houston, TX), and remdesivir (HY-104077) was purchased from MedChemExpress (Monmouth Junction, NJ). Chloroquine was dissolved in Dulbecco’s Phosphate-Buffered Saline (DPBS; Welgene), and all other reagents were dissolved in DMSO for the screening. Anti-SARS-CoV-2 N protein antibody was purchased from Sino Biological Inc. (Beijing, China). Alexa Fluor 488 goat anti-rabbit IgG (H + L) secondary antibody and Hoechst 33342 were purchased from Molecular Probes. Paraformaldehyde (PFA) (32% aqueous solution) and normal goat serum were purchased from Electron Microscopy Sciences (Hatfield, PA) and Vector Laboratories, Inc. (Burlingame, CA), respectively.

### Dose-response curve (DRC) analysis by immunofluorescence

Ten-point DRCs were generated for each drug. Calu-3 cells were seeded at 2.0 × 10^4^ cells per well in EMEM, supplemented with 10% FBS and 1X Antibiotic-Antimycotic solution (Gibco) in black, 384-well, μClear plates (Greiner Bio-One), 24 h prior to the experiment. Ten-point DRCs were generated, with compound concentrations ranging from 0.1-50 μM. For viral infection, plates were transferred into the BSL-3 containment facility and SARS-CoV-2 was added at a multiplicity of infection (MOI) of 0.1. The cells were fixed at 24 hpi with 4% PFA and analyzed by immunofluorescence. The acquired images were analyzed using in-house software to quantify cell numbers and infection ratios, and antiviral activity was normalized to positive (mock) and negative (0.5% DMSO) controls in each assay plate. DRCs were generated in Prism7 (GraphPad) software, with dose-response-inhibition nonlinear regression analysis. IC_50_ and CC_50_ values were obtained with the identical analysis method. Mean values of independent duplicate experiments were used for analysis. Each assay was controlled by Z’-factor and the coefficient of variation in percent (%CV).

## Acknowledgements

The pathogen resource (NCCP43326) for this study was provided by the National Culture Collection for Pathogens. This work was supported by the National Research Foundation of Korea (NRF) grant funded by the Korean government (MSIT) (NRF-2017M3A9G6068245 and NRF-2020M3E9A1041756).

## Notes

### Competing Interest Statement

The authors have declared no competing interest.

## References

1. Zhou P, Yang X-L, Wang X-G, Hu B, Zhang L, Zhang W, Si H-R, Zhu Y, Li B, Huang C-L, Chen H-D, Chen J, Luo Y, Guo H, Jiang R-D, Liu M-Q, Chen Y, Shen X-R, Wang X, Zheng X-S, Zhao K, Chen Q-J, Deng F, Liu L-L, Yan B, Zhan F-X, Wang Y-Y, Xiao G-F, Shi Z-L. 2020. A pneumonia outbreak associated with a new coronavirus of probable bat origin. Nature 579:270–273.

2. Gorbalenya AE, Baker SC, Baric RS, de Groot RJ, Drosten C, Gulyaeva AA, Haagmans BL, Lauber C, Leontovich AM, Neuman BW, Penzar D, Perlman S, Poon LLM, Samborskiy D V., Sidorov IA, Sola I, Ziebuhr J. 2020. The species Severe acute respiratory syndrome-related coronavirus: classifying 2019-nCoV and naming it SARS-CoV-2. Nat Microbiol 5:536–544.

3. Wang M, Cao R, Zhang L, Yang X, Liu J, Xu M, Shi Z, Hu Z, Zhong W, Xiao G. 2020. Remdesivir and chloroquine effectively inhibit the recently emerged novel coronavirus (2019-nCoV) in vitro. Cell Res 30:269–271.

4. Jeon S, Ko M, Lee J, Choi I, Byun SY, Park S, Shum D, Kim S. 2020. Identification of antiviral drug candidates against SARS-CoV-2 from FDA-approved drugs. bioRxiv.

5. Zhu Y, Chidekel A, Shaffer TH. 2010. Cultured Human Airway Epithelial Cells (Calu-3): A Model of Human Respiratory Function, Structure, and Inflammatory Responses. Crit Care Res Pract 2010:1–8.

6. Pruijssers AJ, George AS, Schäfer A, Leist SR, Gralinski LE, Dinnon KH, Yount BL, Agostini ML, Stevens LJ, Chappell JD, Lu X, Hughes TM, Gully KL, Martinez DR, Brown AJ, Graham RL, Perry JK, Pont V Du, Pitts J, Ma B, Babusis D, Murakami E, Feng JY, Bilello JP, Porter DP, Cihlar T, Baric RS, Denison MR, Sheahan TP. 2020. Remdesivir potently inhibits SARS-CoV-2 in human lung cells and chimeric SARS-CoV expressing the SARS-CoV-2 RNA polymerase in mice. bioRxiv.

7. Borba MGS, Val FFA, Sampaio VS, Alexandre MAA, Melo GC, Brito M, Mourão MPG, Brito-Sousa JD, Baía -da-Silva D, Guerra MVF, Hajjar LA, Pinto RC, Balieiro AAS, Pacheco AGF, Santos JDO, Naveca FG, Xavier MS, Siqueira AM, Schwarzbold A, Croda J, Nogueira ML, Romero GAS, Bassat Q, Fontes CJ, Albuquerque BC, Daniel-Ribeiro C-T, Monteiro WM, Lacerda MVG. 2020. Effect of High vs Low Doses of Chloroquine Diphosphate as Adjunctive Therapy for Patients Hospitalized With Severe Acute Respiratory Syndrome Coronavirus 2 (SARS-CoV-2) Infection. JAMA Netw Open 3:e208857.

8. Cao B, Wang Y, Wen D, Liu W, Wang J, Fan G, Ruan L, Song B, Cai Y, Wei M, Li X, Xia J, Chen N, Xiang J, Yu T, Bai T, Xie X, Zhang L, Li C, Yuan Y, Chen H, Li H, Huang H, Tu S, Gong F, Liu Y, Wei Y, Dong C, Zhou F, Gu X, Xu J, Liu Z, Zhang Y, Li H, Shang L, Wang K, Li K, Zhou X, Dong X, Qu Z, Lu S, Hu X, Ruan S, Luo S, Wu J, Peng L, Cheng F, Pan L, Zou J, Jia C, Wang J, Liu X, Wang S, Wu X, Ge Q, He J, Zhan H, Qiu F, Guo L, Huang C, Jaki T, Hayden FG, Horby PW, Zhang D, Wang C. 2020. A Trial of Lopinavir–Ritonavir in Adults Hospitalized with Severe Covid-19. N Engl J Med 382:1787–1799.

9. Hoffmann M, Kleine-Weber H, Schroeder S, Krüger N, Herrler T, Erichsen S, Schiergens TS, Herrler G, Wu N-H, Nitsche A, Müller MA, Drosten C, Pöhlmann S. 2020. SARS-CoV-2 Cell Entry Depends on ACE2 and TMPRSS2 and Is Blocked by a Clinically Proven Protease Inhibitor. Cell 181:271-280.e8.

10. Yamamoto M, Matsuyama S, Li X, Takeda M, Kawaguchi Y, Inoue J, Matsuda Z. 2016. Identification of Nafamostat as a Potent Inhibitor of Middle East Respiratory Syndrome Coronavirus S Protein-Mediated Membrane Fusion Using the Split-Protein-Based Cell-Cell Fusion Assay. Antimicrob Agents Chemother 60:6532–6539.

11. Kawase M, Shirato K, van der Hoek L, Taguchi F, Matsuyama S. 2012. Simultaneous Treatment of Human Bronchial Epithelial Cells with Serine and Cysteine Protease Inhibitors Prevents Severe Acute Respiratory Syndrome Coronavirus Entry. J Virol 86:6537–6545.

12. Klok FA, Kruip MJHA, van der Meer NJM, Arbous MS, Gommers DAMPJ, Kant KM, Kaptein FHJ, van Paassen J, Stals MAM, Huisman MV, Endeman H. 2020. Incidence of thrombotic complications in critically ill ICU patients with COVID-19. Thromb Res.

13. Poissy J, Goutay J, Caplan M, Parmentier E, Duburcq T, Lassalle F, Jeanpierre E, Rauch A, Labreuche J, Susen S. 2020. Pulmonary Embolism in COVID-19 Patients: Awareness of an Increased Prevalence. Circulation CIRCULATIONAHA.120.047430.

